# Multiple RSV strains infecting HEp-2 and A549 cells reveal cell line-dependent differences in resistance to RSV infection

**DOI:** 10.1101/2021.06.15.448622

**Authors:** Anubama Rajan, Felipe-Andrés Piedra, Letisha Aideyan, Trevor McBride, Matthew Robertson, Hannah L. Johnson, Gina Marie Aloisio, David Henke, Cristian Coarfa, Fabio Stossi, Vipin Kumar Menon, Harshavardhan Doddapaneni, Donna Marie Muzny, Sara Joan Javornik Cregeen, Kristi Louise Hoffman, Joseph Petrosino, Richard A Gibbs, Vasanthi Avadhanula, Pedro A. Piedra

## Abstract

Respiratory syncytial virus (RSV) is a leading cause of pediatric acute respiratory infection worldwide. There are currently no approved vaccines or antivirals to combat RSV disease. A few transformed cell lines and two historic strains have been extensively used to study RSV. Here we report a thorough molecular and cell biological characterization of HEp-2 and A549 cells infected with four strains of RSV representing both major subgroups as well as historic and more contemporaneous genotypes -- [RSV/A/Tracy (GA1), RSV/A/Ontario (ON), RSV/B/18537 (GB1), RSV/B/Buenos Aires (BA)] -- via measurements of viral replication kinetics and viral gene expression, immunofluorescence-based imaging of gross cellular morphology and cell-associated RSV, and measurements of host response including transcriptional changes and levels of secreted cytokines and growth factors. Our findings strongly suggest 1) the existence of a conserved difference in gene expression between RSV subgroups A and B; 2) the A549 cell line is a more stringent and natural host of replicating RSV than the HEp-2 cell line; and 3) consistent with previous studies, determining the full effects of viral genetic variation in RSV pathogenesis requires model systems as tractable as transformed cell lines but better representative of the human host.

**IMPORTANCE:** Infection with respiratory syncytial virus (RSV) early in life is essentially guaranteed and can lead to severe disease. In vitro data from two historic RSV/A strains and two cell lines, HEp-2 and A549, constitute most of our knowledge; but RSV contains ample variation from two evolving subgroups (A and B) showing recent convergent evolution. Here we measure viral action and host response in HEp-2 and A549 cells infected with four RSV strains from both subgroups and representing both historic and more contemporaneous strains. We discover a subgroup-dependent difference in viral gene expression and find A549 cells are more potently antiviral and more sensitive, albeit subtly, to viral variation. Our findings reveal important differences between RSV subgroups and two widely used cell lines and provide baseline data for experiments with model systems better representative of natural RSV infection.

## INTRODUCTION

RSV contains considerable genetic diversity with two major subgroups, A and B, each containing multiple genotypes. Subgroups A and B are estimated to have diverged over 300 years ago (1) and show convergent evolution over the last 10-20 years with the emergence of dominant genotypes containing a duplication of less than 100 nucleotides (72 nucleotides in RSV/A, 60 nucleotides in RSV/B) within the G gene (2). Although this duplication appears to result in a modest enhancement of RSV binding strength to host cells (3), its full effects on the viral life cycle and host response are unknown. Furthermore, whole genomes of historic RSV strains (i.e., those isolated over 20 years ago) and more contemporaneous strains from the same subgroup show a sequence divergence of ~5%, which is ~4x lower than that of cognate strains (historic or contemporaneous) from different subgroups. Thus, genetic variation in RSV is substantial in magnitude and interesting in structure. It remains unknown whether the genetic variation separating RSV subgroups and genotypes leads to significant functional differences in the RSV life cycle and whether or to what extent it provokes divergent host responses.

Immortalized respiratory epithelial cell lines, particularly HEp-2 and A549, have been used to investigate the molecular virology and pathogenesis of RSV for decades (4–10). HEp-2 cells were derived from a larynx carcinoma over 60 years ago and show clear evidence of contamination with HeLa cells (11–13); A549 cells were derived from a type II alveolar epithelial carcinoma and have been used to study RSV infection much as HEp-2 cells have. To date, no study has systematically searched for differences between these widely used and seemingly interchangeable cell lines with respect to their permissiveness to RSV infection, their RSV-induced cellular and immune responses, and their sensitivity to different RSV strains.

Here we report a broad and unbiased characterization of in vitro RSV infections using a diverse set of viral strains in two major cell lines. We infected HEp-2 and A549 cells with four RSV strains belonging to both major subgroups and four genotypes representing both historic and more contemporaneous strains: RSV/A/USA/BCM-Tracy/1989 (GA1), RSV/B/WashingtonDC.USA/18537/1962(GB1), RSV/A/USA/BCM813013/2013(ON), RSV/B/USA/BCM80171/2010(BA)). We assessed 1) viral replication kinetics and viral gene expression, 2) the distribution of infecting and/or egressing RSV and accompanying cell morphological changes, 3) host transcriptional changes, and 4) levels of secreted cytokines and growth factors.

Our data indicate that HEp-2 and A549 cells differ systematically in their response to RSV infection, with the latter cell line supporting less viral replication by mounting a more effective antiviral response. Interestingly, facets of the A549 response appear to support the RSV life cycle in ways the HEp-2 response does not; and A549 cells show heightened sensitivity to differences within infecting RSV strains.

## RESULTS

### HEp-2 cells support greater RSV replication than A549 cells

HEp-2 and A549 cells were compared for their ability to support replication of the four RSV strains [RSV/A/Tracy (GA1 genotype), RSV/A/Ontario (ON genotype), RSV/B/18537 (GB1 genotype), RSV/B/Buenos Aires (BA genotype)] (Figure 1). HEp-2 cells showed significantly greater mean levels of intracellular and extracellular viral RNA through time and across strains (Figure 1A and B) [linear regression model, p-value <0.01]. Consistent with viral RNA levels, the amount of infectious virions released was also greater from HEp-2 cells than A549 cells through time and across strains (Figure 1C) [linear regression model, p-value <0.01]. There were no significant differences in growth kinetics (levels of viral RNA or infectious virions) between RSV subgroups or historic [RSV/A/Tracy (GA1) and RSV/B/18537 (GB1)] and contemporaneous ([RSV/A/Ontario (ON) and RSV/B/Buenos Aires (BA)] strains.

**Figure 1:**
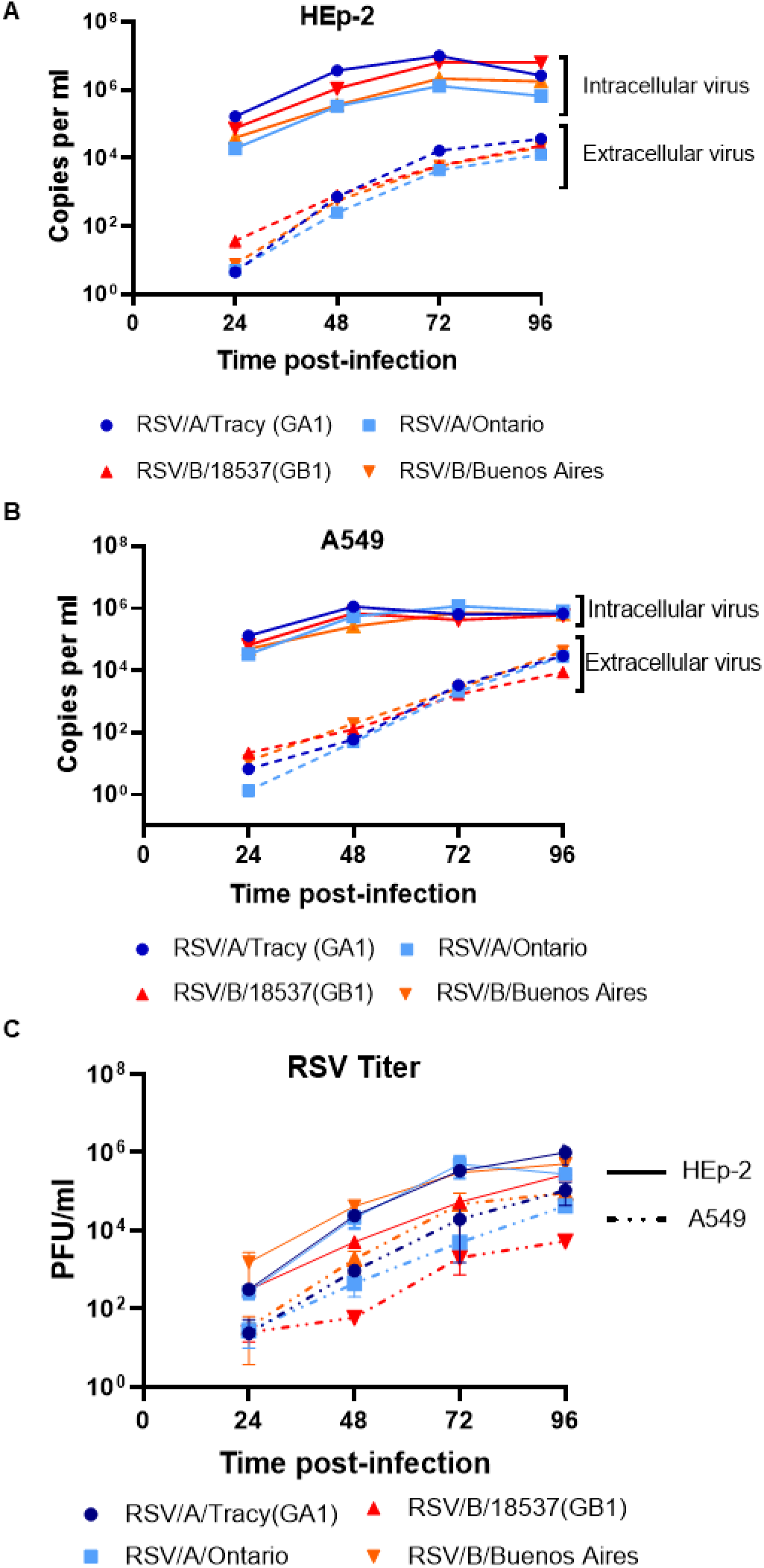
Viral load and replication kinetics of RSV infection. HEp-2 and A549 cells were infected with RSV [RSV/A/Tracy (GA1), RSV/A/Ontario (ON), RSV/B/18537 (GB1), RSVB/Buenos Aires (BA) at an MOI of 0.01. Samples were collected at 24-, 48-, 72- and 96-hour post inoculation. RNA was isolated from media (extracellular) and cell lysate (intracellular) and copy numbers of RSV nucleocapsid (N) gene RNA were determined using quantitative real time PCR (qRT-PCR). Levels of RSV N gene RNA in (A) HEp-2 cells at different time points after RSV inoculation, (B) A549 cells at different time points after RSV inoculation. (C) Extracellular live virus was collected from the media of HEp-2 and A549 inoculated cell cultures. The extracellular virus concentrations were determined by a quantitative plaque assay and reported as log10 plaque forming unit (PFU)/ml in HEp-2 cells. Data shown were from two individual experiments with two replicates per group in each experiment and are represented as mean ± SD.

### Viral gene expression is highly comparable across strains, but transcriptional readthrough shows a major difference between RSV subgroups A and B

We measured genome-wide transcription from RSV strains using high-throughput short read mRNA sequencing (mRNA-seq). Previous measurements by qPCR show that viral gene expression patterns do not vary by cell line (14), so we decided to employ the more widely used HEp-2 cells for the measurements reported here. RSV-infected cell lysate samples were collected for mRNA-seq at time-points (24 and 48 hours post-infection (hpi)) occurring within a previously established window of steady-state gene expression (14), and the resulting coverage plots were averaged to reveal a genome-wide transcription pattern for each infecting strain. Genome-wide transcription patterns were comparable across strains, showing what appear to be three tiers of gene expression (high: NS1-G, medium: F-M2, low: L) (Figure 2A). Percentages of transcriptional readthrough were mostly comparable across strains but showed significant differences at NS2-N, M-SH, and G-F gene junctions (Figure 2B), with the M-SH gene junction showing a large subgroup-dependent difference (Figure 2B). An apparent subgroup-dependent difference in readthrough also occurred at the NS1-NS2 gene junction, but the higher percentage and large error recorded for both subgroup B viruses result from a dip in sequence coverage over the NS1 ORF (data not shown) of unknown origin (Figure 2B).

**Figure 2:**
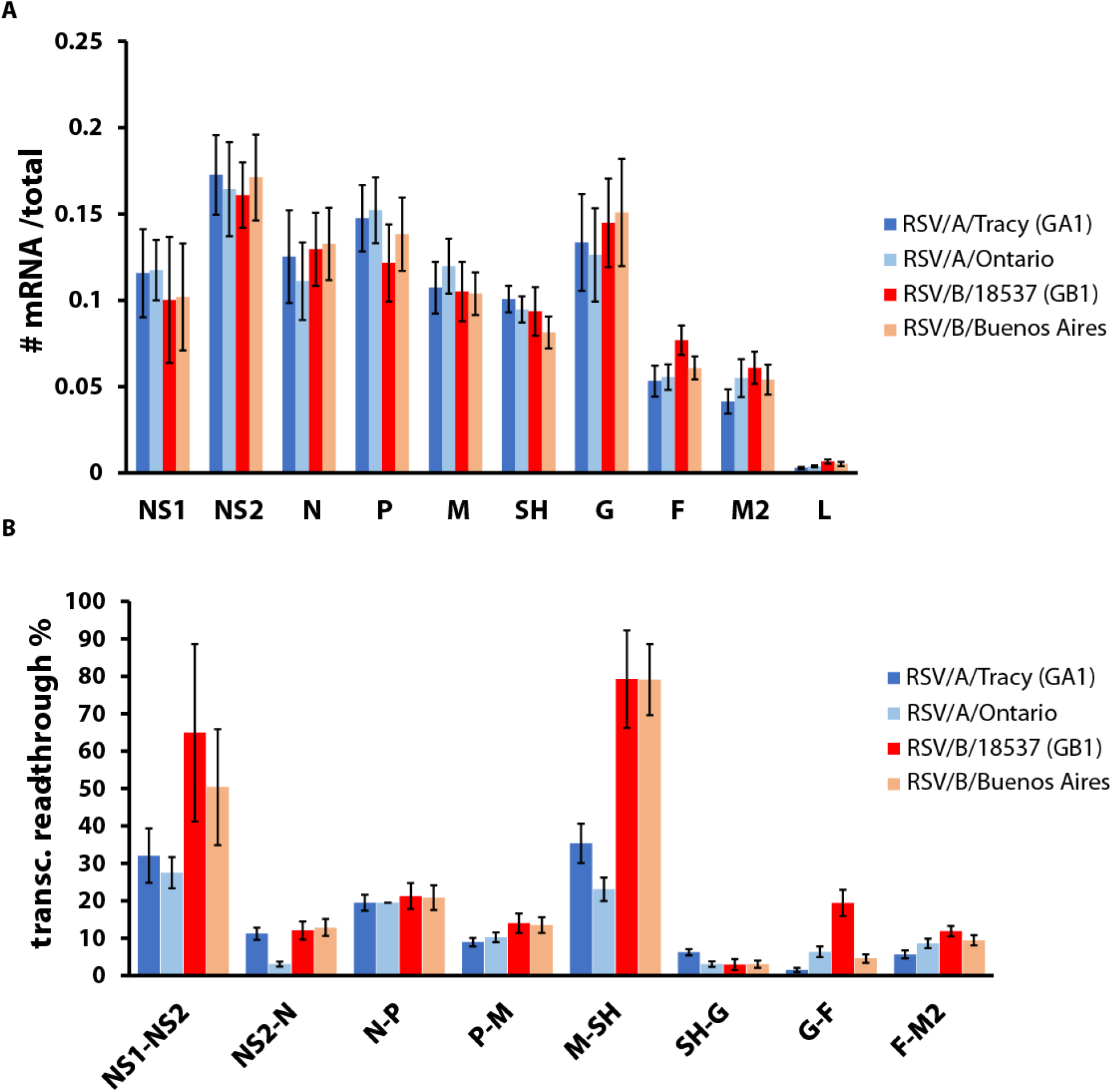
RSV gene expression. HEp-2 cells were infected with RSV [RSV/A/Tracy (GA1), RSV/A/Ontario, RSV/B/18537 (GB1), RSV/B/Buenos Aires] at an multiplicity of infection (MOI) of 0.01. Samples were collected at 24- and 48-hours post-inoculation. RNA was isolated from cell lysates and prepared for and subjected to high-throughput short read sequencing. (A) Relative mRNA levels are comparable across the RSV strains tested. (B) Transcriptional read-through at particular gene junctions varies between RSV subgroups and strains. Data shown are averages ± SD of results for 24- and 48-hours post-inoculation.

### Distribution of RSV and gross morphological changes via rearrangements of the actin cytoskeleton in infected HEp-2 and A549 cells

RSV has been reported to infect small foci of epithelial cells at low multiplicity of infection (MOI) (4, 10). To explore how RSV infects, spreads, and whether it induces changes to cytoskeletal structure in a cell line- and/or RSV strain-dependent way, we infected HEp-2 and A549 cells with RSV strains at low MOI (0.01) and performed epifluorescence deconvolution imaging for actin, cell nuclei, and RSV at 24, 48, 72, and 96 hours post-inoculation (hpi). Regardless of strain, HEp-2 cells showed widespread infection at 24 hpi with RSV levels increasing before reaching a plateau at 72hpi (Figure 3). In contrast, RSV infection in A549 cells was limited to small foci of cells at 24 hpi with subsequent spreading comparable to that seen in HEp-2 cells (Figure 4).

**Figure 3:**
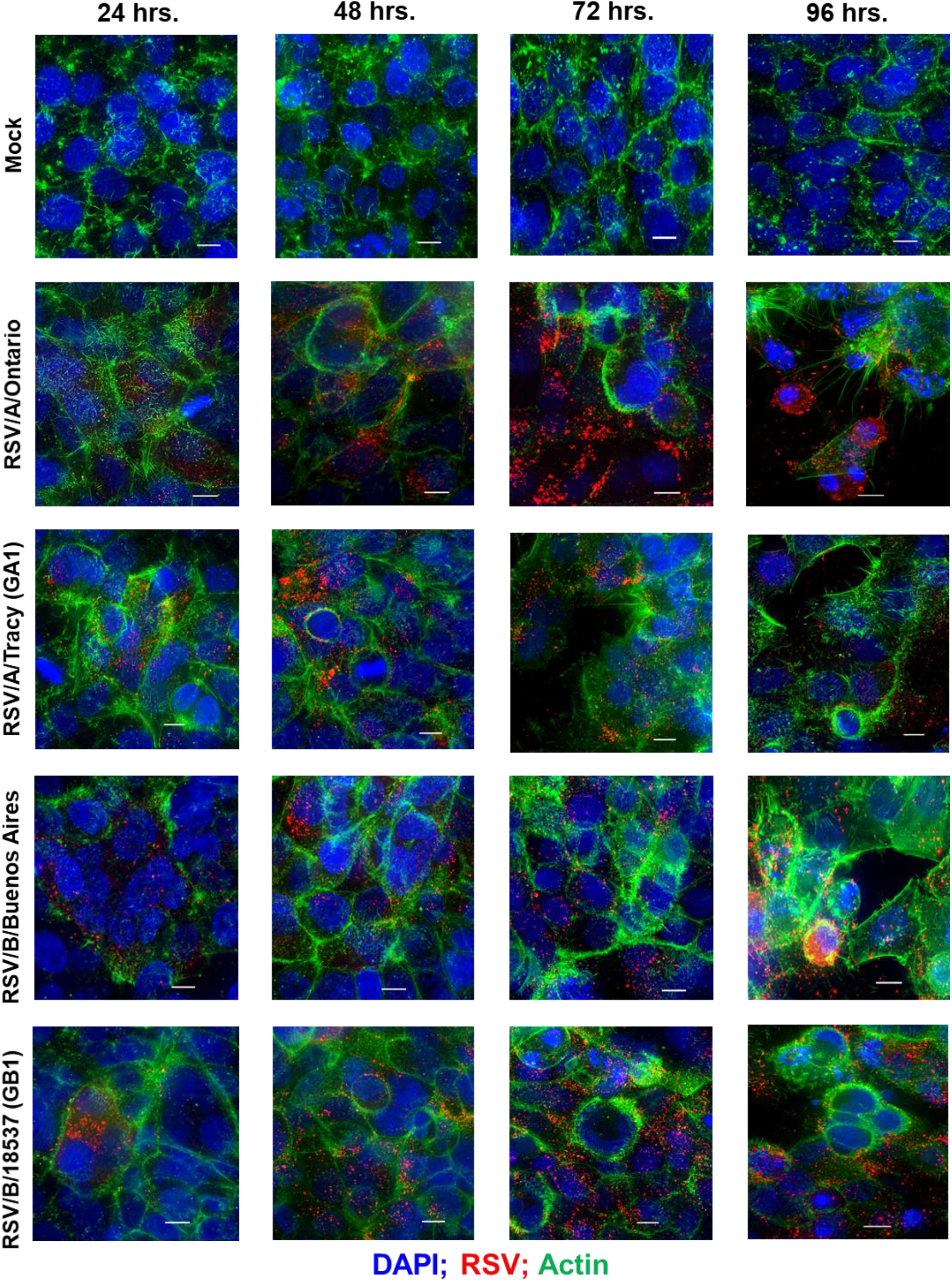
RSV infectivity pattern and cellular damage in HEp-2 cells visualized by immunofluorescence micrographs. Representative epifluorescence deconvolution micrographs of HEp-2 cells labeled or nuclei (DAPI), RSV (Red), and Actin (Green). Cells were either mock treated or infected with RSV/A/Tracy (GA1), RSV/A/Ontario (ON), RSV/B/18537 (GB1) or RSV/B/Buenos Aires (BA) at a multiplicity of infection of 0.01 for 24-, 48-, 72- and 96-hours. Scale bars indicate 10μm.

**Figure 4:**
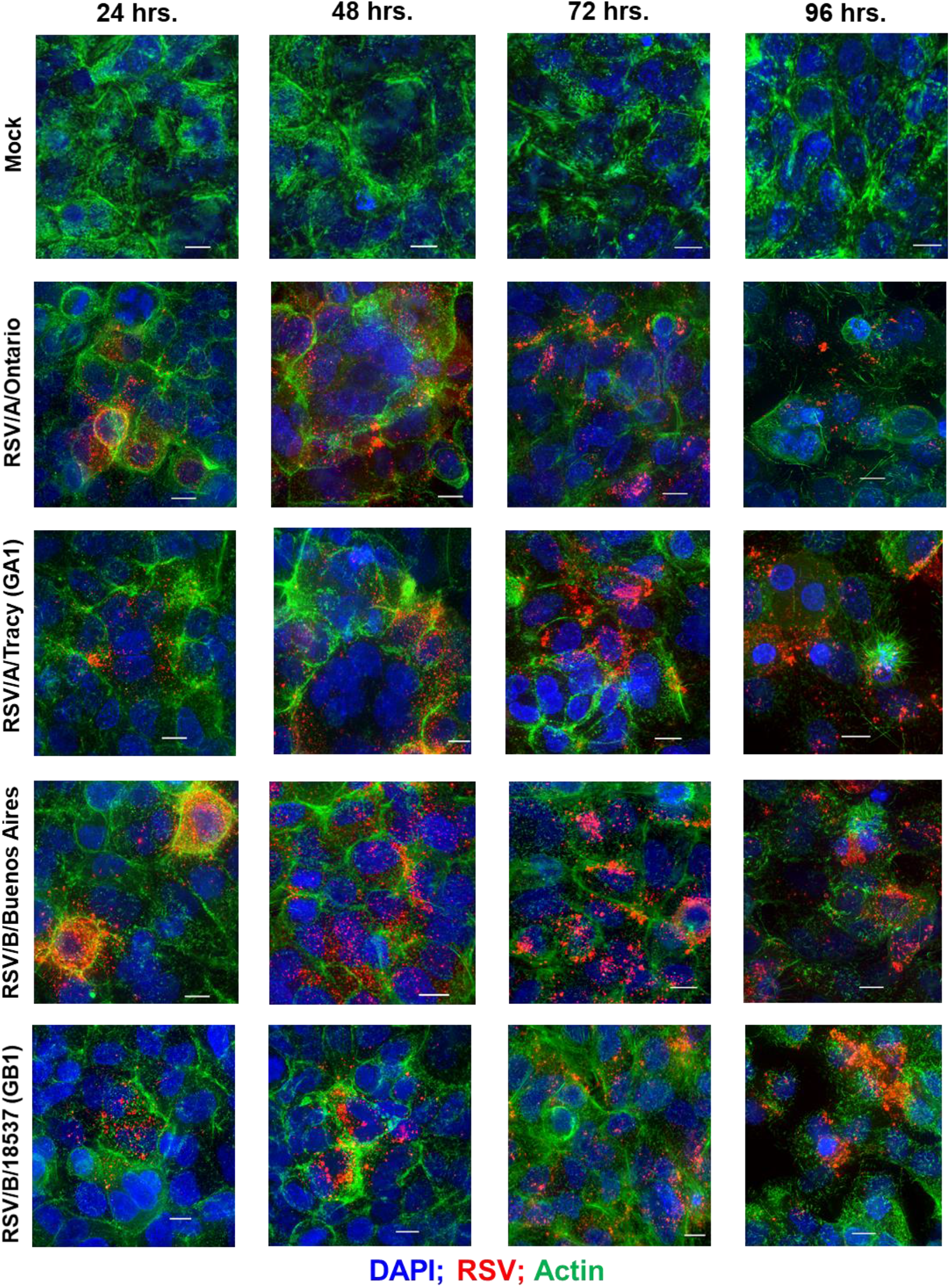
RSV infectivity pattern and cellular damage in A549 cells visualized by immunofluorescence micrographs. Representative epifluorescence deconvolution micrographs of A549 cells labeled for nuclei (DAPI), RSV (Red), and Actin (Green). Cells were either mock treated or infected with RSV/A/Tracy (GA1), RSV/A/Ontario (ON), RSV/B/18537 (GB1) or RSV/B/Buenos Aires (BA) at a multiplicity of infection of 0.01 for 24-, 48-, 72- and 96-hours. Scale bars indicate 10μm.

Actin staining revealed differences in actin rearrangement between infected HEp-2 and A549 cells and differing rearrangements induced by RSV subgroups A and B (Figures 3 and 4). It is worth noting that mock-infected HEp-2 and A549 cells also differed in the morphology of their actin cytoskeleton, with A549 cells containing either a larger number of actin filaments or filament bundles of greater size. Both cell lines show actin rearrangements in response to all four infecting strains at the earliest time-point visualized, 24 hpi, with actin filaments adopting an arrangement reminiscent of thin halos at the boundaries of infected cells. In HEp-2 cells infected with RSVB strains, these actin halos significantly increased in thickness at 72 hpi and beyond. In HEp-2 cells infected with RSVA strains, actin halos rearranged into microns-long tail-like structures at 96 hpi. In addition, RSVA-infected HEp-2 cells show considerable cytopathic effect at 96 hpi, while RSVB-infected HEp-2 cells do not. In contrast, RSV-infected A549 cells displayed actin rearrangements resulting in what appear to be paracellular gaps rather than cellular damage at 96 hpi (Figure 4).

### Global changes in HEp-2 and A549 gene expression in response to RSV infection

We next sought to determine whether the origins of any of the differences observed in RSV infection could be illuminated by interrogating the host transcriptional response. To do so we subjected RSV-infected and mock-infected A549 and HEp2 cells to high-throughput short read RNA sequencing (RNAseq). Principal component analysis (PCA) shows that cell type accounts for most of the variation in the data (Figure 5A), and a strong transcriptional response to RSV in both HEp-2 and A549 cells does not become clear until 72 and 96 hpi regardless of infecting strain (Figure 5A). Of the four time-points assayed, the number of significantly up- and downregulated genes relative to mock-infected cells is low at 24hpi (<100) then rises rapidly beyond 48 hpi before approaching a peak of ~6000 to −8000 genes at 72 hpi and beyond (Figure 5B). Up- and downregulated genes are clustered into a heat map to compare across cell lines, virus strains and time-points (Figure 5C).

**Figure 5:**
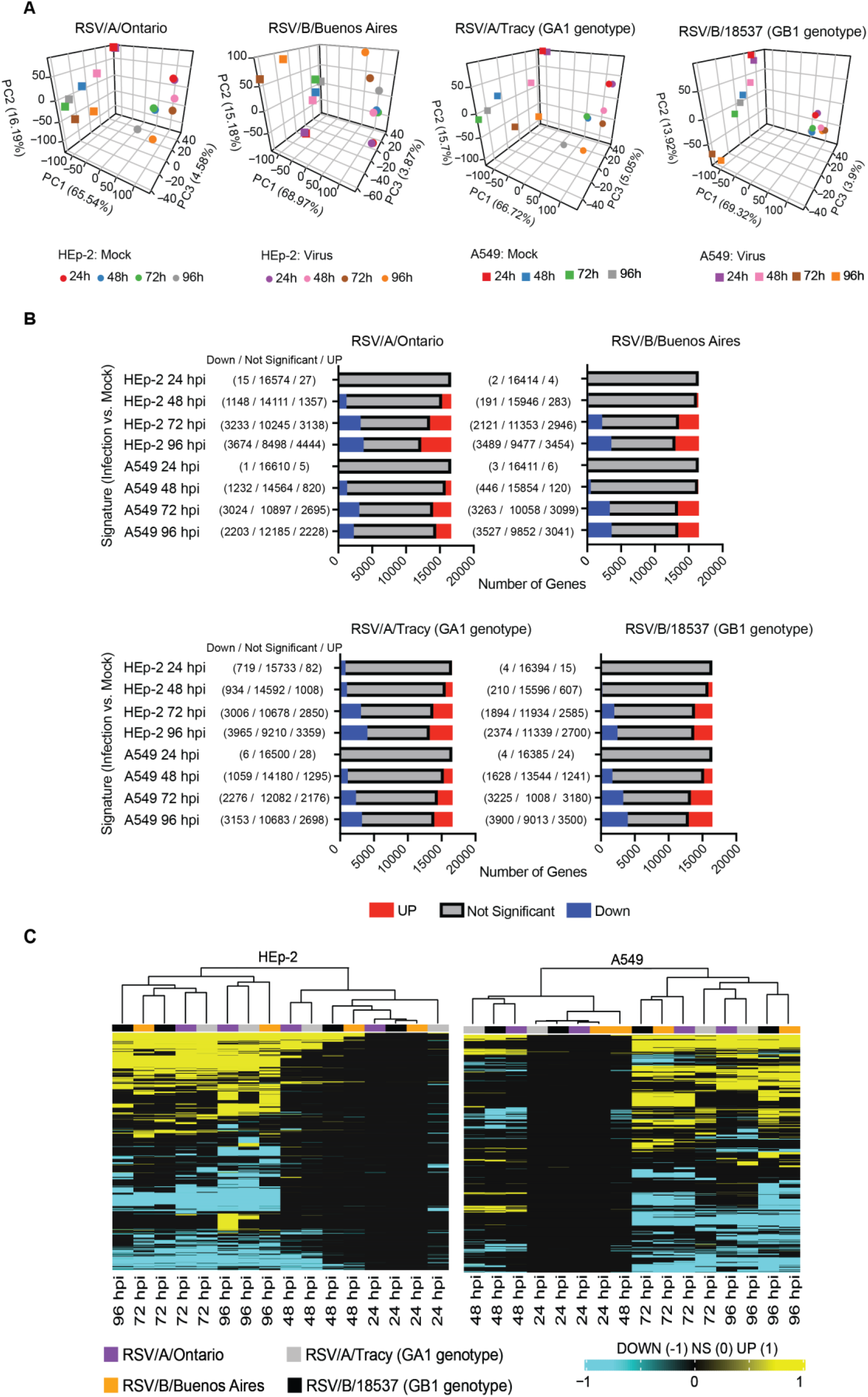
RNA sequencing analysis of RSV infection of HEp-2 and A549 cells (A) Principal component analysis of RSV/A/Ontario (ON), RSV/B/Buenos Aires (BA), RSV/A/Tracy (GA1) and RSV/B/18537 (GB1) infected HEp-2 and A549 cells demonstrating the variability observed in the samples (HEp-2 versus A549 and infected versus mock controls). (B) Total number of significant genes identified for the RSV/A/Ontario, RSV/B/Buenos Aires, RSV/A/Tracy and RSV/B/18537 infected group in HEp-2 and A549 cells (adjusted p-value < 0.05). (C) Samples were hierarchically clustered based on the union of differentially expressed genes across all comparisons (False discovery rate (FDR) <0.05 and fold change exceeding 1.5x). Genes were converted to 1 for up-regulated genes and −1 for down-regulated genes and 0 if it were not significant in a particular signature. Samples were clustered using the Euclidean distance.

### A functional analysis reveals many cell line- and some viral strain-dependent features of the host transcriptional response to RSV infection

The Gene Set Enrichment Analysis (GSEA) software (15) was used to search our data for functional pathways enriched for up- or downregulated genes relative to mock-infected cells (≥ 2x up- or downregulated and false discovery rate (FDR) < 0.05). The pathways identified are mostly related to host immune, cell cycle, and metabolic processes (Figure 6, Supplementary Figures 1–4). Many of the identified functional pathways show normalized enrichment scores (NES) with temporal dynamics that differ strongly between HEp-2 and A549 cells. For instance, the NES for the apoptotic response in A549 cells rises monotonically to a peak at 96hpi regardless of the infecting strain, while the HEp-2 apoptotic response is weaker and dips to near baseline values at 96 hpi (Figure 6, Supplementary Figure 1). Genes involved in progression through the cell cycle (E2F target and G2M checkpoint genes) are upregulated at 48 and 72 hpi in A549 cells, while they are strongly repressed at 72 and 96 hpi in HEp-2 cells (Supplementary Figure 2). Genes upregulated in response to interferon alpha (IFN-α) and interferon gamma (IFN-γ) are activated beyond 24 hpi in A549 cells, while they are strongly repressed at 24 hpi in HEp2 cells with the NES gradually rising to <0.5 at 96 hpi for all four RSV strains assayed (Figure 6, Supplementary Figure 3). Genes encoding components of the inflammasome appear highly upregulated during at least one time-point for HEp-2 cells infected with either of the four RSV strains, while their expression state does not ever differ significantly from mock in A549 cells (Supplementary Figure 3). The NES for genes involved in cytokine signaling rises monotonically to a peak of 1-1.5 at 96 hpi in A549 cells but plateaus at lower values between 48 and 72 hpi in HEp-2 cells, with RSV/A/Tracy-infected HEp-2 cells being the only combination showing modest upregulation (NES ~0.9) through both 48 and 72 hpi in HEp-2 cells (Supplementary Figure 3). Genes upregulated in response to increases in IL2 and IL6 via STAT5 and STAT3, respectively, show mostly switch-like but modest activation between 48 and 72 hpi in A549 cells, while showing gradual and comparably modest activation plateauing beyond 72 hpi in HEp-2 cells (Figure 6, Supplementary Figure 4). The NES for genes involved in the inflammatory response gradually rises through the course of the experiment in A549 cells, with the two RSV/A infections showing significant peak upregulation (NES 1.8-1.9) at 96 hpi; in contrast, the NES plateaus at more modest values beyond 48 hpi in infected HEp-2 cells (Supplementary Figure 4). Genes upregulated by NFκB in response to TNFα have an NES that increases in switch-like fashion beyond 48 hpi in A549 cells, rising to modest activation (NES 0.8-1) at 96 hpi; the NES for these genes is higher at and plateaus beyond 48 hpi in infected HEp-2 cells (Supplementary Figure 4).

**Figure 6:**
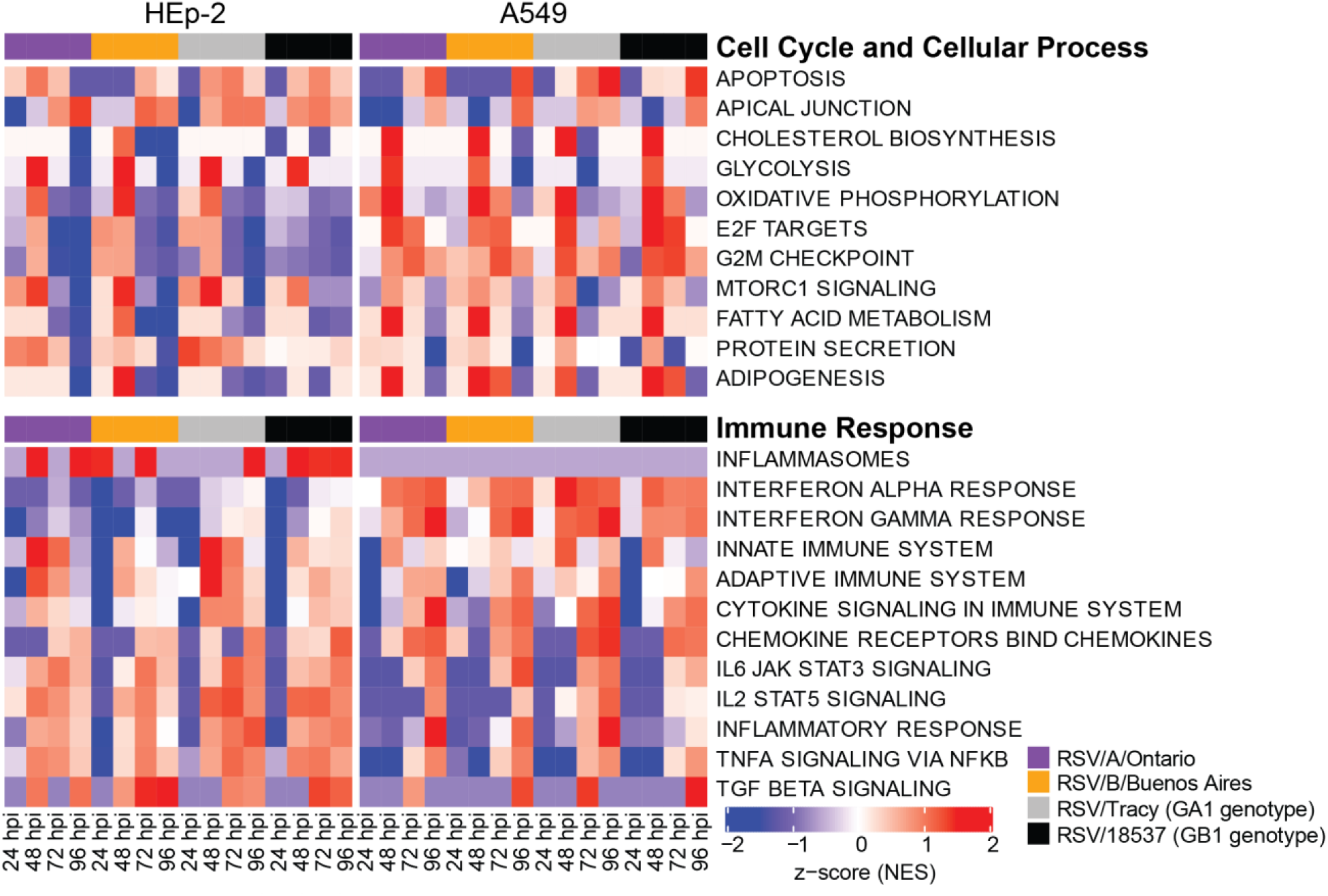
Select reactome and hallmark gene ontology pathways identified from a Gene Set Enrichment Analysis by filtering for the top ten normalized enrichment scores (NES) for each category. False discovery rate of 0.05 was used in the pathway filtering.

The two cell lines also show highly similar responses to RSV in a few pathways (Figure 6, Supplementary Figures 1 and 2). Both show 1) gradual activation of genes encoding components of the apical junction complex; 2) significant upregulation of glycolysis and oxidative phosphorylation at 48 hpi with two minor exceptions showing no upregulation at that time-point (glycolysis: RSV/A/Tracy:A549; oxidative phosphorylation: RSV/B/18537:HEp-2); and 3) gradually increasing activation of chemokine receptor genes through 96 hpi.

Some metabolic pathways even show a host response in apparent favor of the viral life cycle: genes associated with cholesterol biosynthesis, fatty acid metabolism, and adipogenesis are highly upregulated regardless of infecting strain in A549 cells at 48 hpi; a similar but lower peak in upregulation occurs in HEp-2 cells but only in response to the RSV/B/Buenos Aires strain (Figure 6, Supplementary Figures 1 and 4)

We also observed a few instances of apparent RSV subgroup-dependent and historical-vs.-contemporaneous-strain-dependent differences in host response, especially in A549 cells (Supplementary Figures 2 and 3). In response to the two RSV/A strains assayed, genes upregulated in response to TGF-β1 are strongly activated at 72 hpi before reverting to a repressed state at 96hpi; whereas in response to the two RSV/B strains assayed these genes remain repressed until 96 hpi when they switch to being strongly upregulated. Both the expression status of genes upregulated by STAT5 in response to IL2 stimulation and genes involved in protein secretion show an apparent dependence in A549 cells on whether the infecting RSV strain is historic or more contemporaneous. The latter also seems to hold in HEp2 cells; in both cell lines, genes associated with protein secretion are strongly downregulated at 96 hpi only when the infecting strain is contemporaneous (RSV/A/Ontario or RSV/B/Buenos Aires).

### Cytokine profiles of HEp-2 and A549 cells in response to RSV infection

Finally, we probed the host response to RSV infection using a Luminex™ platform to measure levels of 29 cytokines and growth factors in cell media. 23 different cytokines and growth factors were detected in response to RSV infection, all with levels showing strong differences between HEp-2 and A549 cells. Interleukin (IL)-6 showed some of the highest increases relative to mock in both HEp-2 and A549 cells regardless of the infecting strain at 72 and 96 hpi (Figure 7). IL-8 levels showed comparable increases at later time-points in infected HEp-2 cells but, with the exception of infections with RSV/A/Tracy, were greatly reduced in infected A549 cells (Figure 7). Matrix metalloproteinase-9 (MMP-9), fibroblast growth factor-2 (FGF-2), granulocyte-macrophage colony-stimulating factor (GM-CSF), and tumor necrosis factor-α (TNF-α) showed much smaller increases than IL6 and IL8 but were the next highest detected in infected HEp-2 cells across RSV strains (Figure 7A and 7B). Regulated upon Activation, Normal T Cell Expressed and Presumably Secreted (RANTES) showed increases comparable to or exceeding IL6 in infected A549 cells (Figure 7C and 7D). Infected A549 cells also showed significant increases in C-X-C motif chemokine ligand 11 (CXCL11) at 96 hpi from RSVA strains, and large increases in G-CSF and interferon gamma-induced protein 10 (IP-10) at 96 hpi from the more contemporaneous strains RSV/A/Ontario and RSV/B/Buenos Aires.

**Figure 7:**
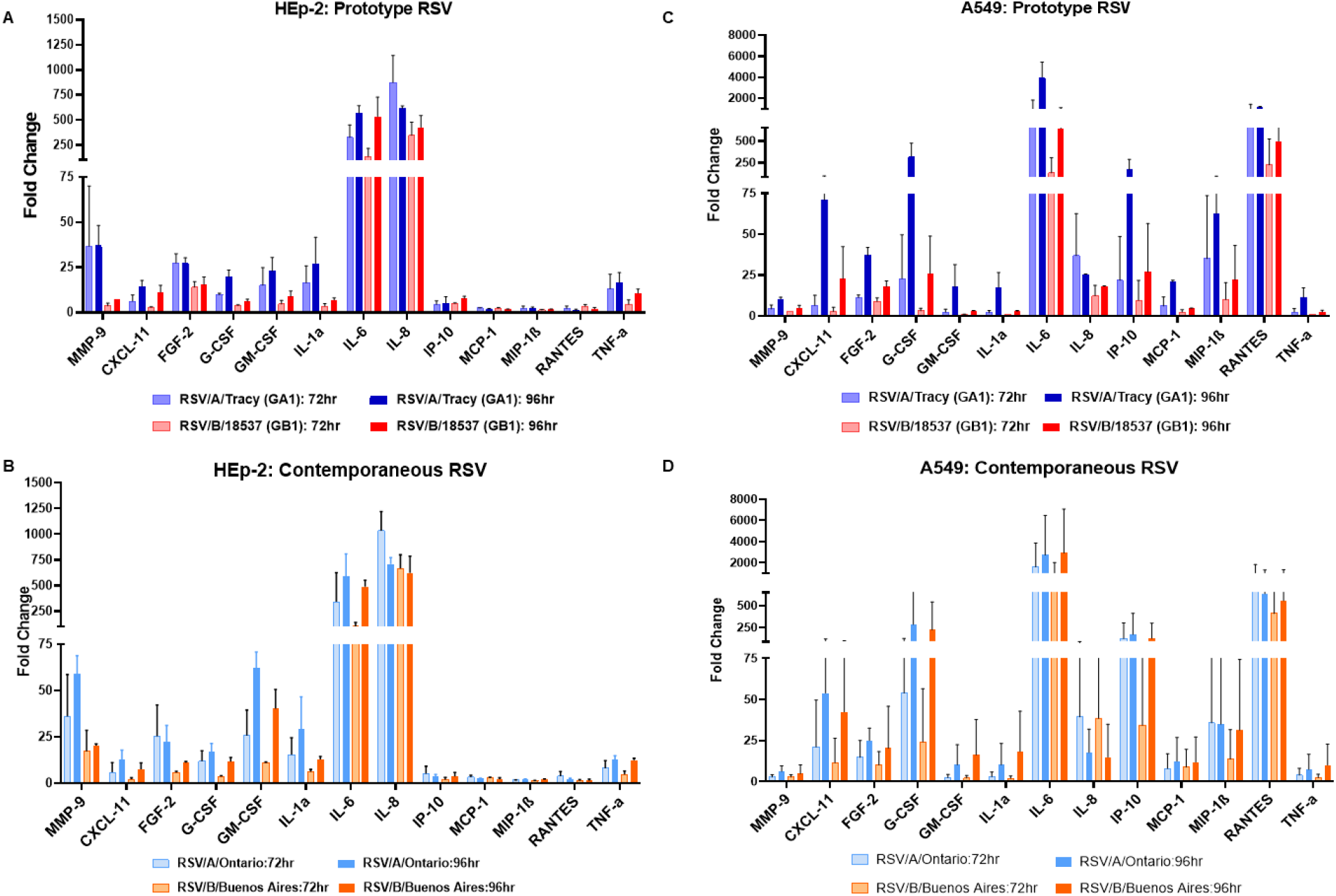
Profiles of releasing cytokines and chemokines from HEp-2 and A549 cells. Cells were infected with RSV/A/Tracy (GA1) or RSV/B/18537 (GB1) or RSV/A/Ontario (ON) or RSV/B/Buenos Aires at a multiplicity of infection of 0.01. The cultured supernatants were harvested from mock-infected or RSV-infected cells at 24-, 48-, 72-, 96-hour post-inoculation (hpi). Profiles of extracellular cytokines and chemokines released in the supernatants were determined by the Multiplex-Luminex cytokine assay. The fold changes of each cytokine or chemokine between virus- and mock-infected cells were shown. Fold changes of cytokines and chemokines released from HEp-2 cells infected with (A) prototype RSV strains (RSV/A/Tracy (GA1) and RSV/B/18537 (GB1)) and (B) contemporaneous RSV strains (RSV/A/Ontario (ON) and RSV/B/Buenos Aires. Fold changes of cytokines and chemokines released from A549 cells infected with prototype RSV strains (C) and contemporaneous RSV (D) strains. At least two independent experiments were performed, and the ratios were presented as mean ± SD.

Similar to the cytokine kinetics observed in RSV-infected HEp-2 and A549 cells, the maximum fold increase in lactate dehydrogenase (LDH) and caspase 3/7 levels relative to mock occurred at 72 and 96 hpi (Figure 8). The relative fold increase in caspase 3/7 was comparable between infected HEp-2 and A549 cells (Figure 8A and 8B), while the relative fold increase in LDH was approximately two-fold higher in HEp-2 cells than A549 cells (Figure 8C and 8D).

**Figure 8:**
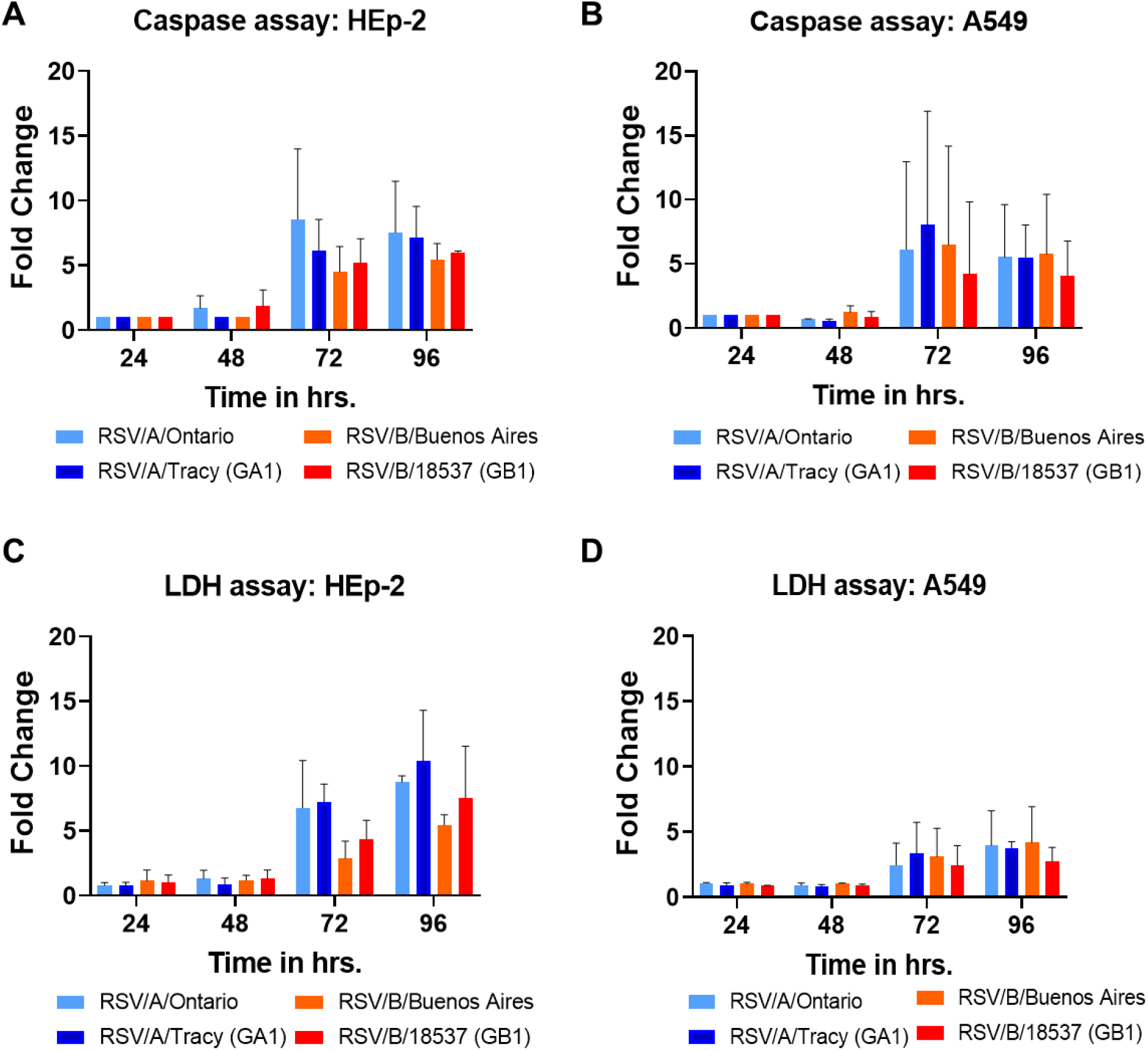
Lactate dehydrogenase (LDH) and caspase assay on HEp-2 and A549 cells infected with RSV strains. Cells were infected with RSV/A/Tracy (GA1) or RSV/B/18537 (GB1) or RSV/A/Ontario (ON) or RSV/B/Buenos Aires at a multiplicity of infection) of 0.01. The cultured supernatants were harvested from mock-infected or RSV-infected cells at 24-, 48-, 72-, 96-hour post-inoculation. Fold changes of caspase released from (A) HEp-2 cells and (B) A549 cells are shown. Fold changes of LDH released from (C) HEp-2 cells and (D) A549 cells are shown. At least two independent experiments were performed, and the ratios were presented as mean ± SD.

## DISCUSSION

We completed a broad and thorough characterization of RSV infections in vitro, discovering a subtle but potentially conserved difference in gene expression between RSV subgroups A and B, numerous differences in host response to RSV infection between two widely used cell lines (HEp-2 and A549), and subtle but varied evidence of host cell line sensitivity to RSV genetic variation, especially between subgroups and between historical (no duplication in G gene) and more contemporaneous (duplication in G gene) strains.

Measurements of viral replication kinetics revealed HEp-2 cells to be more permissive to RSV infection than A549 cells, giving rise to ~10-fold more infectious virus at each of four time-points collected over 96 hours. Perhaps consistent with their greater permissibility, confocal imaging revealed greater cytopathic effect in RSV-infected HEp-2 than A549 cells at later time-points (72 and 96 hpi). The two cell lines show global transcriptional responses to RSV infection that are similar in magnitude and kinetics, but with a set of illuminating differences in the regulation of genes involved in several metabolic, immune, and cellular processes. In all, A549 cells gave rise to a more potently antiviral state than HEp-2 cells, with much greater activation of genes upregulated by IFN-α and IFN-γ. The HEp-2 response to RSV infection is more pro-inflammatory in nature. HEp-2 cells show activation of genes involved in inflammasome production and earlier and more sustained activation of genes involved in the inflammatory response. In addition, HEp-2 cells produce elevated levels of both IL6, IL8 and LDH in response to infection with either of the four RSV strains assayed, while A549 cells show elevated levels of both only in response to RSV/A/Tracy. Cytokine profiles for infected A549 cells also show elevated levels of chemo-attractants like RANTES, IP-10, and MIP-1β in addition to increases in IL6. Also, RSV-infected A549 cells showed higher ratios of caspase 3/7 to LDH levels (relative to mock-infected cells) compared to RSV-infected HEp-2 cells. This is consistent with A549 cells manifesting a more potent antiviral (versus proinflammatory) response than HEp-2 cells upon infection with RSV. Levitz et al. observed significantly increased levels of both IL6 and RANTES in A549 cells at 24 hpi in response to an RSV/B strain (NH1125) lacking the G gene duplication that is characteristic of RSV/B/Buenos Aires strains versus an RSV/B strain (NH1067) with the G gene duplication (16). We did not observe such a difference between RSV/B/18537 (lacking G gene duplication) and the RSV/B/Buenos Aires (containing G gene duplication) strains used here. However, the difference observed by Levitz et al. decreases with decreasing MOI, becoming smallest at the lowest MOI (0.013) used (at 24 hpi, levels of both cytokines measured by both groups are quite low -- mostly well below a maximum of ~50 picogram/ml). Our experiments were performed at a still lower MOI (0.01), chosen to maximize the length of our experiment by delaying the onset of significant cytopathic effect. Reverting to a general comparison of the two cell lines, it is interesting that A549 cells showed evidence of a transcriptional response favorable to the RSV life cycle: at 48 hpi, and for all infecting strains, genes involved in cholesterol biosynthesis, fatty acid metabolism, and adipogenesis are strongly activated. This upregulation might involve the production of additional host membrane in support of the generation of new virions; it might also support the production of additional cholesterol-rich rafts promoting further infection through enhanced virus binding to host (17) and also release of new virus particles (18). Features of a host response that promote some aspects of the viral life cycle are not inconsistent with a broader host response that is more potently antiviral and ultimately reduces the permissibility of A549 cells to RSV infection. Continuing the comparison of the two cell lines, A549 cells also showed more instances of strain-dependence in their RSV response, suggesting heightened sensitivity to variation in infecting RSV strains. A549 cells also displayed an apoptotic response that rose monotonically to a peak at 96 hpi, while the HEp-2 apoptotic response was weaker and plateaued before dipping to a magnitude near baseline (except in RSV/A/Ontario-infected HEp-2 cells which showed strong repression of the apoptotic response at 96 hpi), suggesting a less coordinated and effective response from HEp-2 cells at resisting RSV infection. Finally, it is worth considering the origins of these cell lines: A549 cells are derived from a type II alveolar epithelial carcinoma, while the HEp-2 cells that have been in use for decades (the line was originally derived over half a century ago from a larynx carcinoma) are widely known to have resulted from HeLa contamination (11–13). Thus, A549 cells would seem to be the more natural host of RSV infection. Moreover, the combination of an enhanced antiviral response plus some element of a response apparently supporting RSV infection and the heightened sensitivity to RSV variation observed here, all suggest RSV-infected A549 cells to be the model that is better representative of natural RSV infection (i.e., RSV infection of whole human hosts).

A recent study similar to ours compared the host response to a single strain of RSV in A549 and BEAS-2B, a virus-transformed bronchial epithelial cell line, finding that BEAS-2B were less permissive to RSV infection and mounted a more potently antiviral response than the more pro-inflammatory response of A549 cells (4). These results are consistent with those reported here, but because the authors used a single strain of RSV, BEAS-2B sensitivity to variation within RSV could not be assessed. If BEAS-2B cells are better representative of natural RSV infection (as A549 cells are versus HEp-2 cells) we would predict a comparable if not increased level of sensitivity to variation within RSV. We would also expect features of the BEAS-2B response to be in apparent support of the RSV life cycle, as clearly RSV has evolved to infect the respiratory epithelia.

Within each cell line, each of the four RSV strains, with the exception of RSV/B/18537 at all time-points but 24 hpi, replicated with comparable efficiency. However, cell lines did show some subtle sensitivity to infecting strain (mostly at subgroup and historic-vs.-contemporaneous level) at the gross morphological level and in their transcriptional response and production of cytokines.

Unlike most of the strain-dependent differences reported here, the most striking difference observed in cell morphology occurred in HEp-2 cells: RSV/A-infected HEp-2 cells showed actin protrusions several microns in length extending from the cell surface at 96 hpi; and RSV/B-infected HEp-2 cells showed highly dense networks of actin filaments at the same time-point. The origins of these morphological differences, both between RSV subgroups and the two cell lines, are unclear.

Transcriptional data and cytokine levels revealed a few instances of host sensitivity, mostly in A549 cells, to the infecting RSV strain. Genes upregulated in response to TGF-β1 showed a response that appeared strongly RSV subgroup-dependent in A549 cells. In response to RSV/A infections, these genes were repressed until 72hpi when they were strongly activated; in response to RSVB infections, these genes remained repressed until 96hpi when they were strongly activated. This difference seemed largely due to differential regulation of the SMAD7 gene (data not shown). Both the expression status of genes upregulated by STAT5 in response to IL2 stimulation and genes involved in protein secretion showed an apparent dependence in A549 cells on whether the infecting RSV strain is historic (without G gene duplication) or more contemporaneous (with G gene duplication). The latter also seems to hold in HEp2 cells; in both cell lines, genes associated with protein secretion were strongly repressed at 96 hpi only when the infecting strain was contemporaneous. It is tempting to speculate that the duplication within the G gene of both strains (the G or attachment protein is expressed in both an integral membrane (IM) and a truncated non-IM form) leads to a similar host response with respect to the post-translational processing of the G protein, potentially through the shared introduction of new glycosylation sites. Furthermore, production of the chemoattractant CXCL11 appeared subgroup-dependent in A549 cells, with RSV/A strains inducing higher levels of the chemokine than RSV/B strains. Mean levels of the pro-inflammatory cytokine IL6, the maturational cytokine G-CSF, and the chemoattractant IP-10 were over 5-fold higher in response to contemporaneous vs. historic RSV strains at 96 hpi in A549 cells, with much smaller differences if any at 72 hpi (with the exception of IL6 in response to RSV/A/Ontario (ON)) and earlier. These subtly different host responses highlight our lack of knowledge concerning the functional effects of well-known differences within RSV.

Measurements of viral gene expression revealed a single major difference in patterns of gene expression between subgroups A and B, with the two RSV/B strains supporting high transcriptional readthrough at the M-SH gene junction. This is predicted to result in lower expression of SH protein in cells infected with RSV/B strains due to increased production of dicistronic transcripts containing the SH ORF in a position distal to the 5’ cap (19, 20). This appears consistent with published phylogenetic trees suggesting weaker co-evolution between RSV/B SH and the other RSV surface proteins (G and F) than RSV/A SH and G and F (21). Because the function of SH protein remains mysterious, it is not clear how this difference might relate to the few instances of a subgroup-dependent host response reported here. Another apparent subgroup-dependent difference in transcriptional readthrough occurred at the NS1-NS2 gene junction, but the large error recorded for both subgroup B viruses resulted from a dip in sequence coverage over the NS1 ORF (data not shown) of unknown origin. This conserved dip might result from the presence of a cryptic gene end (GE) signal. Regardless, the dip in sequence coverage over the NS1 ORF artificially increases the calculated readthrough percentage at the NS1-NS2 gene junction for the two subgroup B strains assayed.

In brief summary, our thorough characterization of in vitro RSV infections identified a number of viral and host-related features worth further study. These data reveal important differences in host response to RSV infection between two widely used continuous cell lines and provide a baseline for in vitro studies using more realistic model systems of RSV infection such as human airway organoids (10).

## MATERIALS AND METHODS

### Cell culture

HEp-2 and A549 cells were cultured in Minimum Essential Medium (MEM) (Corning 10-010-CM), supplemented with 10% fetal bovine serum (Hyclone SH30070.03), 1% of 10000 U/ml penicillin/streptomycin/ 25μg/mL Fungizone™ (Gibco 15240062), 1 % of L-Glutamine (200 mM) (Gibco 25030081) and maintained in 5% CO_2_ at 36 °C.

### RSV infection, PCR, and plaque assays

Nearly confluent HEp-2 and A549 cells were infected with RSV/A/USA/BCM-Tracy/1989 (GA1), RSV/B/WashingtonDC.USA/18537/1962 (GB1), RSV/A/USA/BCM813013/2013(ON), RSV/B/USA/BCM80171/2010 (BA) at a multiplicity of infection 0.01 for 1.5 hours and inoculum was removed, washed with PBS, and 2% FBS/MEM was added and incubated for a period of 24-, 48-, 72- and 96 hpi. Viral RNA was detected using RT-PCR with primers targeting the nucleocapsid gene of RSV as previously described (22). Virus titer was calculated using semi-quantitative plaque assay as previously described (23).

#### Statistical analysis of RSV replication kinetics data

Separate linear regression models were made to test log-transformed 1) intracellular viral RNA levels, 2) extracellular viral RNA levels, and 3) levels of infectious virions (= plaque forming units) for relationships with the following covariables: cell type, time, and RSV strain. A backwards selection of covariables was conducted in each of the three analyses. Analyses were performed using R version 4.0.3.

### RNA sequencing analysis

#### RNA isolation, Library Preparation, construction, and Sequencing

RSV infections were performed on HEp-2 and A549 cell lines as described above and infected for 24-, 48-, 72- and 96 hours. After infection, host cells were lysed using Trizol and RNA was extracted using Mini Viral RNA Kit (Cat. No. / ID: 52904; Qiagen Sciences, Germantown, Maryland) and automated platform QIAcube (Qiagen, Hilden, Germany) according to the manufacturer instructions (22). All processing steps were performed by Alkek Center for Metagenomics and Microbiome Research, Baylor College of Medicine, Houston, TX, USA. mRNA was enriched using oligo (dT) beads, followed by cDNA synthesis using random hexamers and reverse transcriptase. The final cDNA library was created after purification, terminal repair, ligation of sequencing adapters, size selection, and PCR enrichment.

#### RNA-sequencing analysis, quality control

Fastq files were trimmed to remove adapters and base-pairs that did not meet standards using Trim Galore (Version 0.4.1) (24, 25). Sequences were aligned against the human genome hg38 (GRCh38) using Hisat2 (version 2.1.0), sorted through samtools (version 1.5) (26, 27) and count matrix was generated by mapping reads using featureCounts (version 1.6.0) (28). Non-overlapping uniquely mapped features were counted for further analysis.

#### Expression quantification and gene set enrichment analysis

Differential gene expression (DGE) was determined using the R-packages edgeR, voom following published guidelines (15, 29). Count matrix was filtered for the coding genes, and a cutoff of 1 count per million (CPM) was used to remove low expression genes. The filtered counts were normalized using the trimmed mean of M-values (TMM). Genes were ranked by according to their log2 fold gene expression for use in gene set enrichment analysis (GSEA version 3.0) and analysis was performed using ranked gene lists against MSigDB database (version 6.1) (15, 29).

### Multiplex Luminex cytokine analysis

Cytokine and chemokine levels from HEp-2 and A549 cell supernatant were determined using multiple Milliplex cytokine/chemokine magnetic bead panel (Millipore) according to the manufacturer’s instructions. Samples were assayed on the Magpix (Millipore) using xPonent software (Luminex). The kits used in this study include 1) Milliplex Human Cytokine Panel with Eotaxin/CCL11, FGF-2, G-CSF, GM-CSF, IL-1a, IL-1b, IL-6, IL-8/CXCL8, IL-17E/IL-25, IP-10/CXCL10, MCP-1, MCP-3, MIG, MIP1a, MIP1b, RANTES/CCL5, TNFa, VEGF-A, IL-33, TRAIL, TSLP, TAC/CXCL11, IL-29, BAFF and HMGB1, 2) TGFb1 Singleplex kit, 3) Milliplex Human MMP Panel 2 with MMP9 and MMP7 and 4) Milliplex Human TIMP Panel 2 with TIMP1.

### Lactate dehydrogenase (LDH) and caspase 3/7 analysis

LDH and caspase 3/7 were measured in the supernatant of RSV infected HEp-2 and A549 cells as previously described (30). In brief, total LDH activity was measured in the supernatant using the LDH Cytotoxic Detection Kit Plus, (Roche Applied Science, Indianapolis, IN, USA) following protocol instructions. L-lactate dehydrogenase (Roche Applied Science) was used to construct a standard curve that demonstrated an ample linear dynamic range (r=0.998) at the dilutions tested from 3.9 to 125 milli-units per milliliter (mU/mL). Caspase 3/7, a marker of apoptosis, was measured using the Caspase-Glo-3/7 kit ((Promega, Madison, WI, USA). The luminescence assay was measured using Biotek Synergy H1 microplate reader (Biotek) and expressed as relative luminescence units (RLU). Purified Caspase 3 from ENZO (CAT BML-SE169) was used as the standard for Caspase Glo 3/7 assays and Gen5 Imager Software was used for analysis.

### Immunofluorescence staining and High-Resolution Imaging

Immunofluorescence staining was performed to identify changes in actin filament rearrangement after RSV infection in HEp-2 and A549 cells. HEp-2 and A549 cells were infected with RSV strains as described above on 96 well ibidi μ plates (ibidi, Germany Cat.No:89626). The plates were fixed with 4% paraformaldehyde and incubated at room temperature (RT) for 15 minutes at 24-, 48-, 72- and 96 hpi. The cells were permeabilized and blocked with 1% bovine serum albumin (BSA) with 0.1% Triton X-100 diluted in PBS for 30 min at RT. RSV was detected using anti-RSV antibody from Abcam (Catalog number: ab20745) at 1:1000 dilution in 1% BSA overnight incubation at 4°C. Nuclei and actin were stained with 4′, 6′-diamidino-2-phenylindole (DAPI) (1ug/ml) and Alexa Fluor™ 568 Phalloidin (Catalog number: A12380) at 1:1000 dilution, respectively, for 15 minutes at RT. High resolution automated imaging was performed using a Cytiva DV Live epifluorescence image restoration microscope using an Olympus PlanApo N 60X/1.42 NA objective and a 1.9kx1.9k pco.EDGEsCMOS 5.5 camera with a 1024×1024 FOV. The filter sets used were DAPI (390/18 excitation, 435/48 emission), TRITC (542/27 excitation, 594/45 emission) and Cy5 (632/22 excitation, 676/34 emission). Z stacks (0.2um) covering the whole cell (~10um) were acquired before applying a conservative restorative algorithm for quantitative image deconvolution using SoftWorx v7.0 and saving images as max pixel intensity projections.

---

**Supplementary Figure 1:**
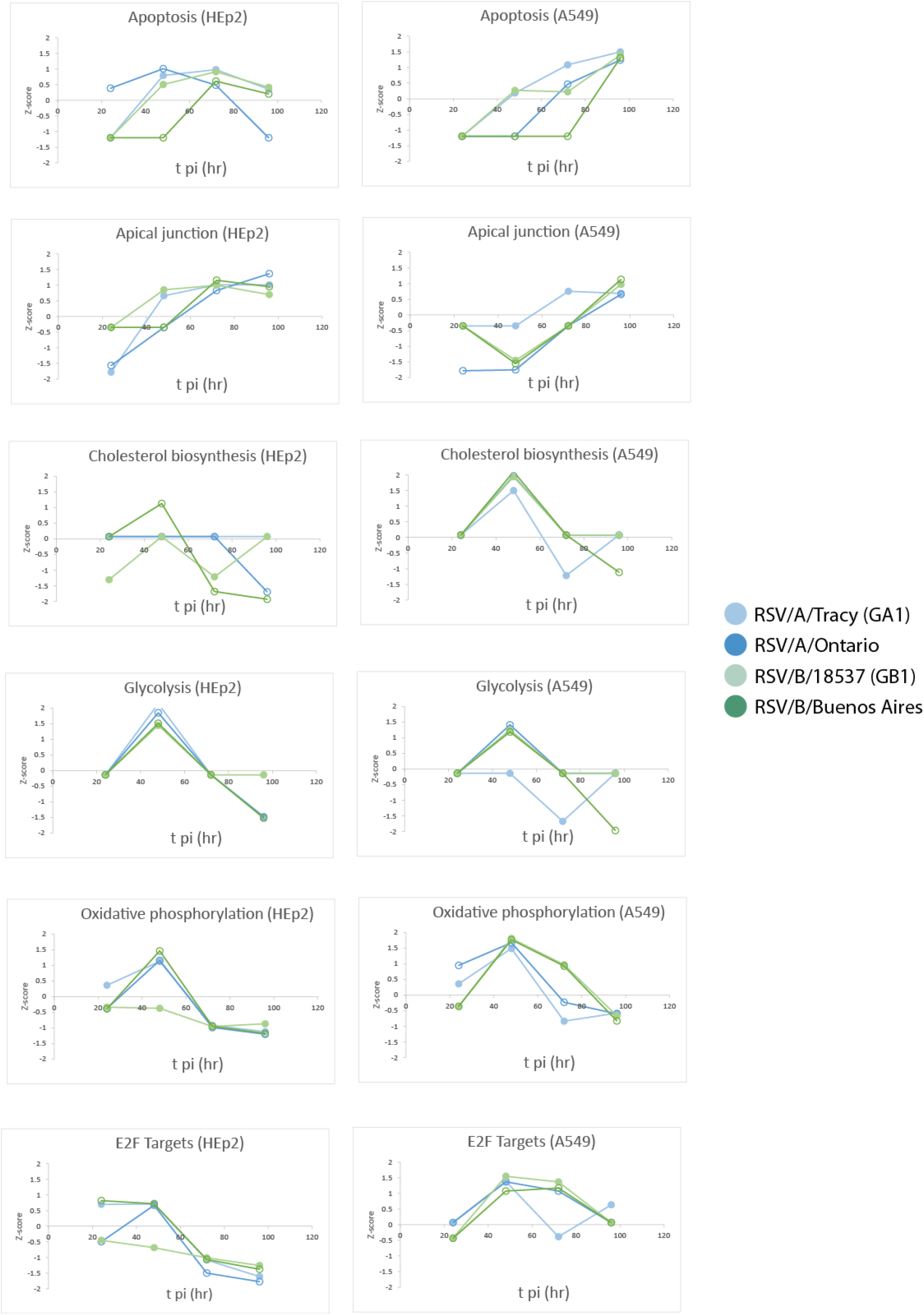
Normalized enrichment scores (NES) through time for Gene Set Enrichment Analysis (GSEA) identified cellular and metabolic pathways enriched for up- or downregulated genes relative to mock.

**Supplementary Figure 2:**
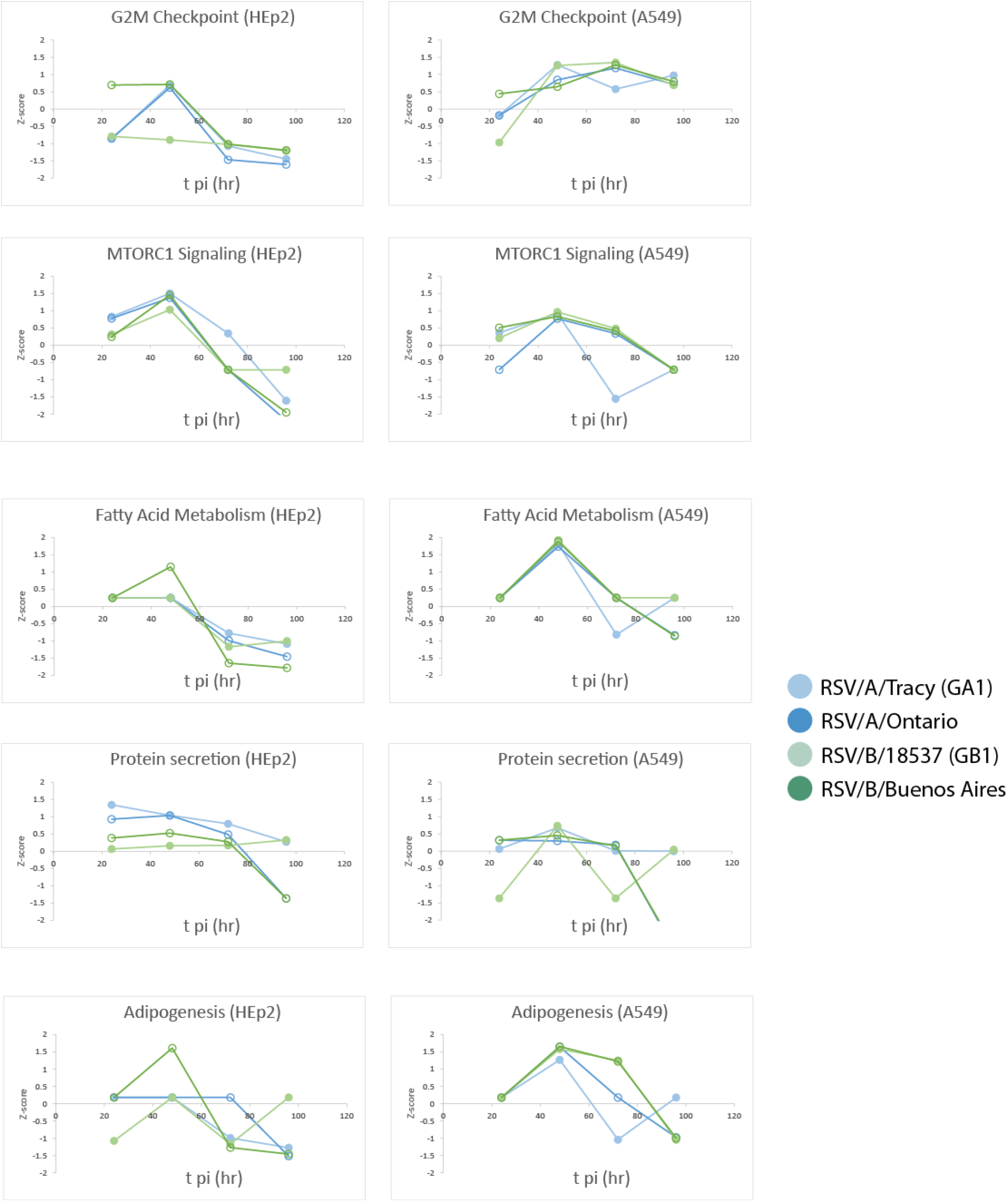
(Continued from Supp. Fig. 1) Normalized enrichment scores (NES) through time for Gene Set Enrichment Analysis (GSEA) identified cellular and metabolic pathways enriched for up- or downregulated genes relative to mock.

**Supplementary Figure 3:**
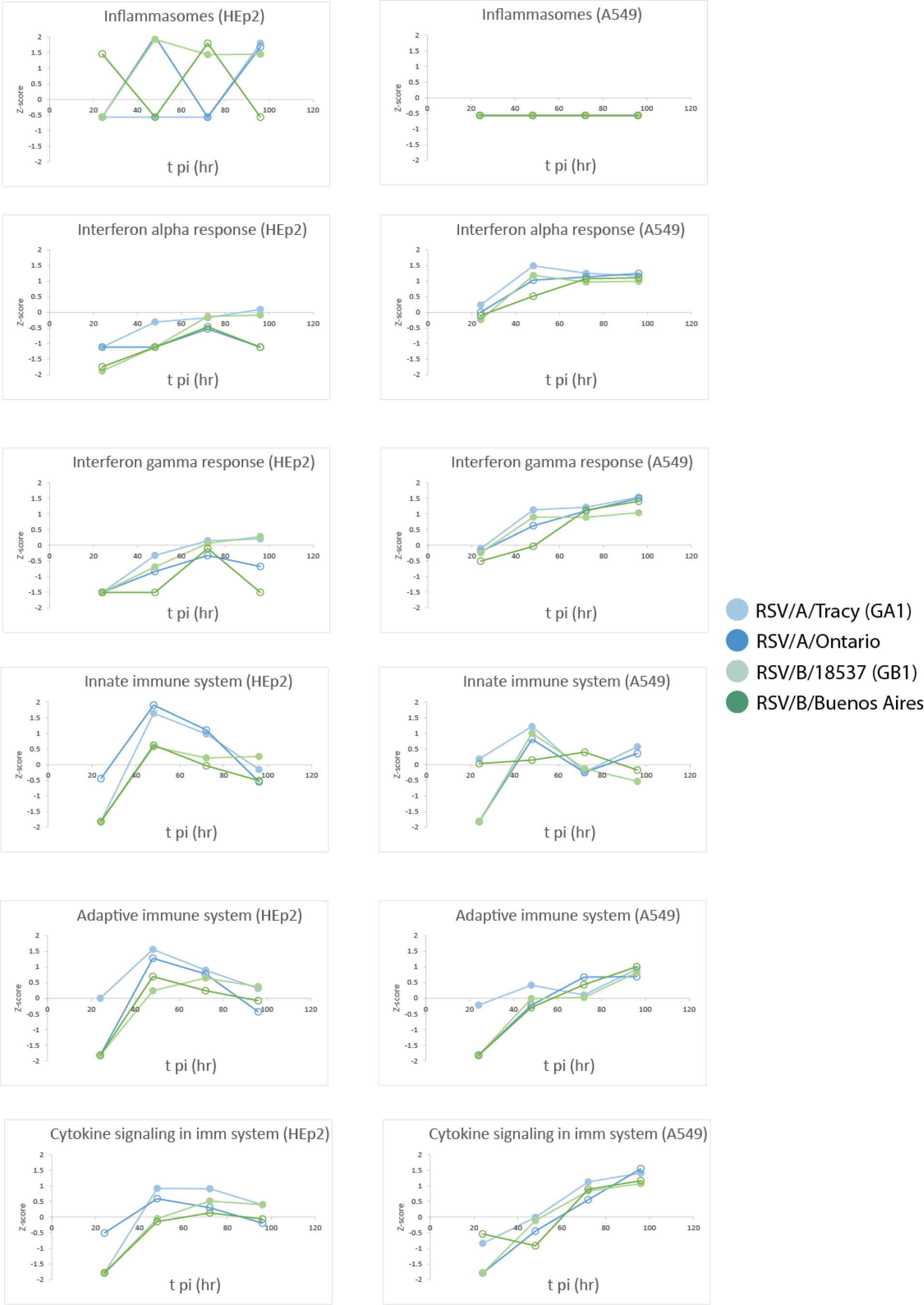
Normalized enrichment scores (NES) through time for Gene Set Enrichment Analysis (GSEA) identified immune pathways enriched for up- or downregulated genes relative to mock.

**Supplementary Figure 4:**
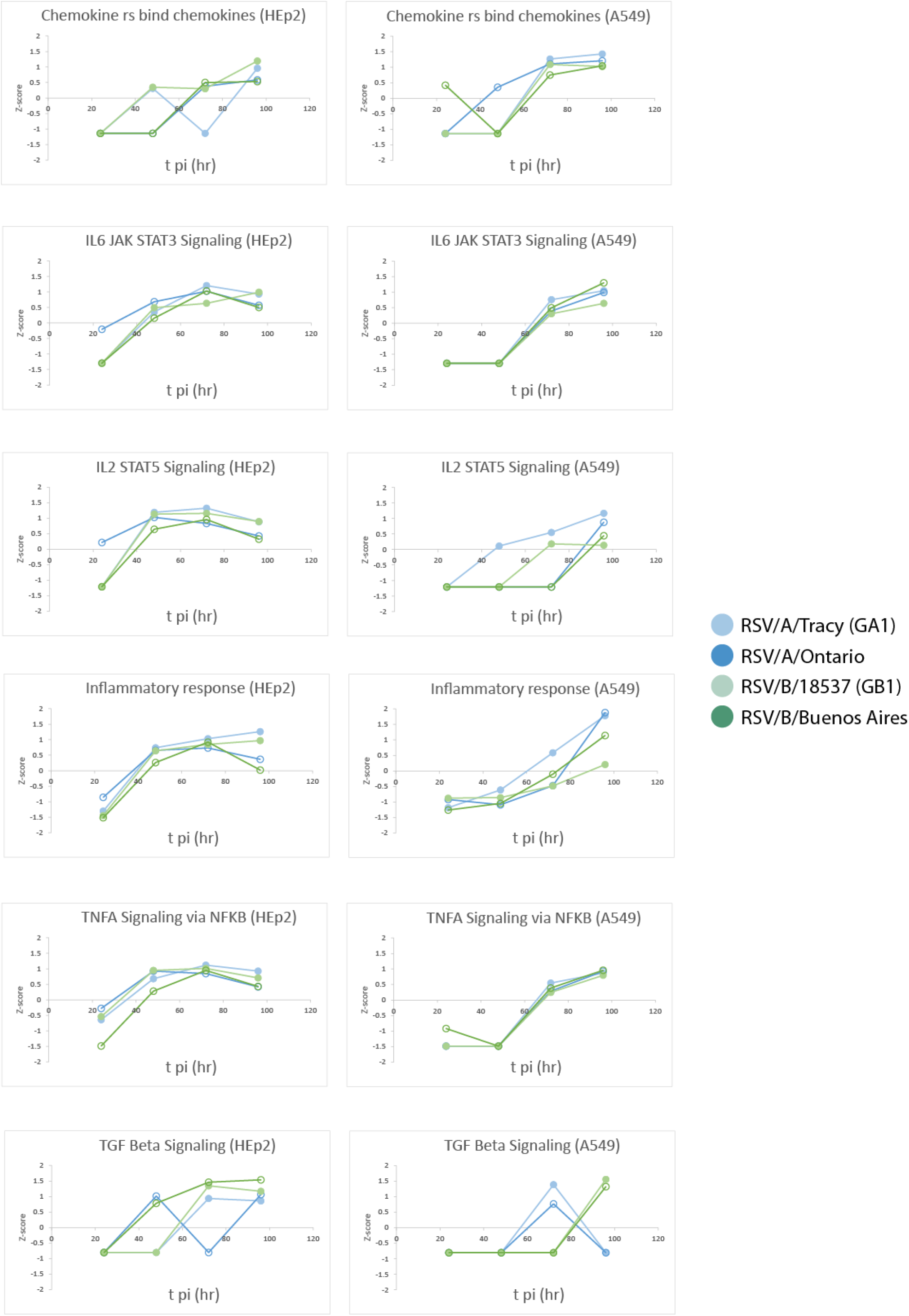
(Continued from Supp. Fig. 3) Normalized enrichment scores (NES) through time for Gene Set Enrichment Analysis (GSEA) identified immune pathways enriched for up- or downregulated genes relative to mock.

